# Spheroid culture remodels mitosis and the proteome in tumor cells

**DOI:** 10.64898/2026.03.12.711371

**Authors:** Ana Petelinec, Claudia Cavarischia-Rega, Adrian Perhat, Boris Maček, Iva M. Tolić

**Affiliations:** Division of Molecular Biology, Ruđer Bošković Institute, Zagreb, Croatia; Quantitative Proteomics, Department of Biology, Interfaculty Institute of Cell Biology, University of Tübingen, Germany

## Abstract

Mitosis depends on precise spindle assembly and positioning, processes influenced by cell shape, size, and microenvironment. Most mechanistic insights into mitosis come from two-dimensional (2D) monolayer cultures, which lack the spatial constraints and extracellular matrix found in tissues, leaving the influence of the tissue environment on mitosis poorly understood. Here, we combine high-resolution imaging and quantitative proteomics to compare mitosis in three-dimensional (3D) multicellular spheroids, generated by magnetic levitation, with that in 2D monolayers. Using a non-transformed cell line and three cancer cell lines from breast, bone, and ovary, we show that 3D culture reshapes mitotic cells and their spindles. Tumor spheroids exhibited a prometaphase delay together with minor chromosome alignment defects, yet chromosome segregation remained largely accurate. Cells in spheroids were rounder, and their spindles were smaller, with increased multipolarity and defects in orientation and position, which varied by cell line. Proteomic profiling revealed broad downregulation of mitotic regulators in spheroids, including kinesins (KIF11, KIF4A), spindle checkpoint proteins, and APC/C components, accompanied by enrichment of metabolic and mitochondrial pathways. Together, our results reveal both shared and cell line-specific modes of mitotic restructuring and establish a framework that connects proteome state to mitotic architecture in 3D environments.

## Introduction

Mitosis ensures faithful genome transmission by assembling a bipolar spindle that aligns and segregates chromosomes with precision (Pavin & Tolić, 2016; Valdez et al., 2023). Spindle assembly, geometry, and positioning are tightly coupled to cell shape, size, and the surrounding microenvironment (Fink et al., 2011; Jiang, 2015; Lázaro-Diéguez et al., 2015). Our current understanding of spindle mechanics, checkpoint signaling, and mitotic errors in healthy and tumor cells has been shaped mostly by studies in two-dimensional (2D) monolayer cultures, where cells grow on flat, rigid substrates that promote spreading. While technically convenient, especially for high-resolution microscopy, monolayers fail to capture essential features of tissues, including confinement, extracellular matrix (ECM) deposition, spatial heterogeneity, and multicellular architecture (Abbas et al., 2023; Da Silva André & Labouesse, 2024; Kapałczyńska et al., 2018).

Three-dimensional (3D) culture systems address many of the limitations inherent to 2D models (Ćosić & Petelinec, 2025; Kapałczyńska et al., 2018; Marques et al., 2022). Because mitosis is highly sensitive to mechanical forces and spatial constraints, culture dimensionality is expected to influence the fidelity and organization of the mitotic machinery. Indeed, several studies have reported that physical confinement, which can occur during 3D growth, perturbs spindle dynamics and cytokinesis (Cattin et al., 2015; Charnley et al., 2013; Desmaison et al., 2013, 2018; M. A. Lancaster et al., 2013). However, other work has shown that 3D growth within a tissue promotes mitotic fidelity, whereas cells dissociated and cultured in 2D exhibit increased chromosome segregation errors (Knouse et al., 2018). These findings reveal an apparent paradox in which dimensionality and cellular context can either compromise or safeguard mitotic accuracy, depending on the biological setting.

Beyond structural differences, global proteomic analyses have identified substantial differences between 2D and 3D cultures (Kim et al., 2018; Kumar et al., 2008). Yet, comparative analyses that couple structural changes with proteomic profiling across cell lines with matched 2D–3D samples are missing, hindering mechanistic insight into mitotic remodeling in a 3D context.

The most physiologically relevant 3D systems, including ex vivo tumor slices, organoids, and in vivo imaging approaches, preserve critical aspects of tissue organization, but their availability can be limited, and comparisons with matched 2D controls are not straightforward (Drost & Clevers, 2018; Hsieh et al., 2025; Marconi et al., 2025). To enable direct 2D–3D comparisons, cell lines that grow efficiently as monolayers can be adapted to form multicellular spheroids by using low-adhesion plates, hanging drops, hydrogel micro-molds, or magnetic levitation (Malhão et al., 2022; Shri et al., 2017; Souza et al., 2010). In the magnetic levitation approach, cells are incubated with magnetic nanoparticles and exposed to an external magnetic field that lifts them to the air–medium interface, where they rapidly aggregate into spheroids (Souza et al., 2010). This scaffold-free method promotes the assembly of spheroids with native ECM, which are flattened, facilitating imaging (Haisler et al., 2013; Souza et al., 2010). Importantly, magnetic levitation does not impair cell proliferation, as cell numbers in spheroids and monolayers increase at similar rates during the first four days of culture, after which spheroids continue to proliferate exponentially for up to nine days, whereas monolayers show only linear growth (Souza et al., 2010). The magnetic force on the cells was estimated to be very low, around 10 pN, sufficient to lift the cells off the dish bottom but too weak to impose significant mechanical stress (Souza et al., 2010).

Here, we generate multicellular spheroids via magnetic levitation and directly compare mitosis in these 3D structures with that in conventional 2D monolayers, using a non-transformed cell line and three cancer cell lines of distinct tissue origins: breast, bone, and ovary. By integrating high-resolution imaging of mitosis with quantitative proteomics, we uncover that 3D growth affects mitotic timing and spindle architecture and position. These changes are accompanied by altered expression of mitotic kinesins and regulatory proteins, including KIF11, CDC20, BUB1B, Cyclin A2, KIF4A, and NuMa. Our findings reveal both shared and cell-line-dependent ways in which mitosis is restructured in 3D and how proteomic remodeling contributes to the altered mitotic behavior.

## Results

### Spheroids and monolayers show similar mitotic fractions, but different interphase morphologies

To compare mitosis in 2D versus 3D culture, we cultured cells in monolayers and as spheroids, using a non-tumor cell line (RPE1 p53KD) and three cancer cell lines of different tissue origin: MDA-MB-231 (breast), U2OS (bone), and OVSAHO (ovary). Spheroids were generated using the magnetic cell levitation method (Souza et al., 2010) (Figure 1A, Methods). These spheroids displayed an irregular, oblate morphology, with pronounced flattening along the z-axis (Figure 1B), consistent with previous observations in other cell types (Haisler et al., 2013; Souza et al., 2010). The longest dimension of the spheroids in the xy-plane was 1–2 mm, whereas their height along the z-axis was tens of times smaller, about 20–50 µm, equivalent to roughly 2–4 cell layers.

**Figure 1.**
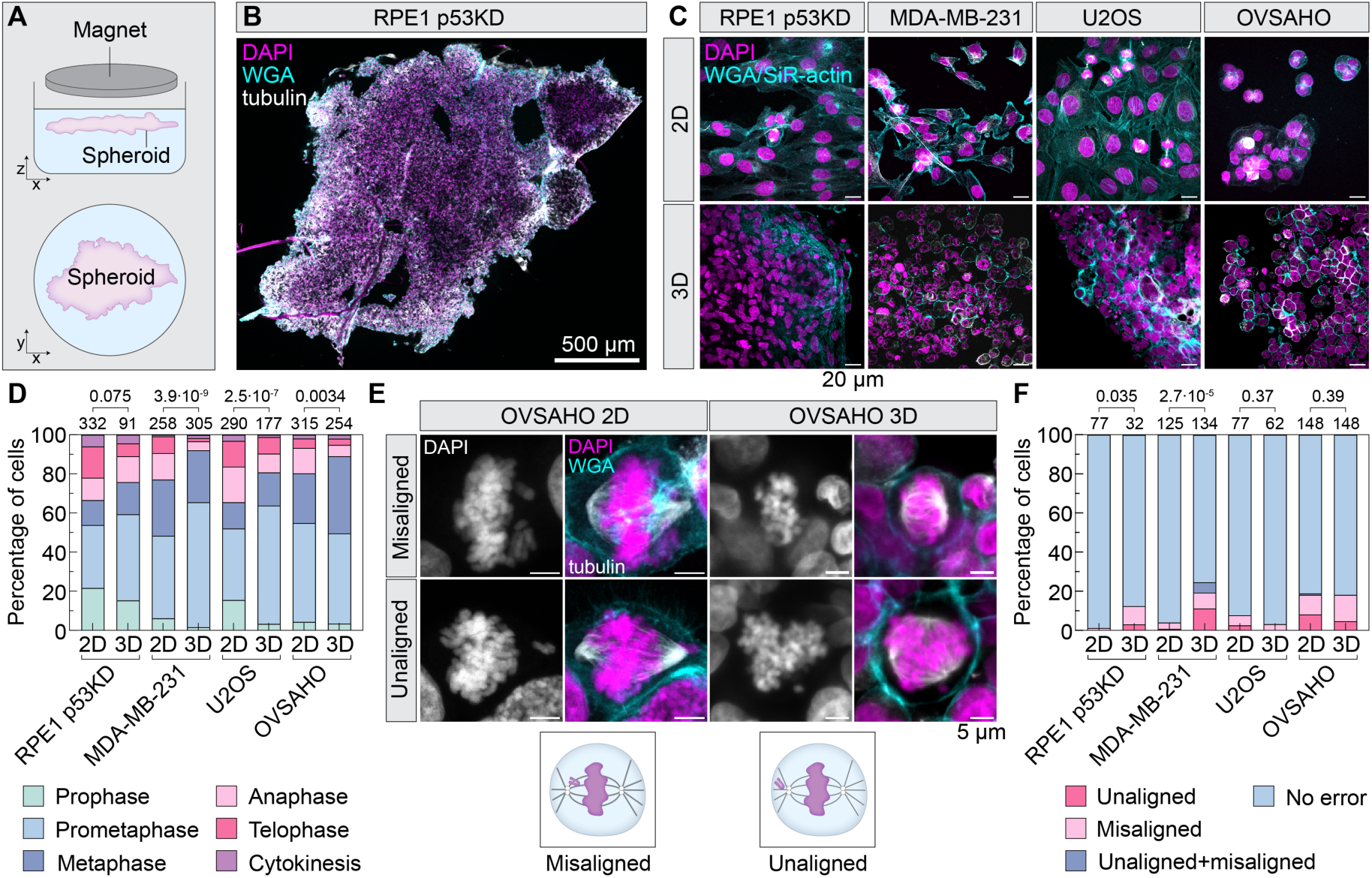
Cells grown in spheroids show a prometaphase delay and minor chromosome misalignment. (A) Schematic view of the magnetic levitation method. (B) A multicellular spheroid of RPE1 p53KD cells stained with DAPI to label DNA (magenta), WGA to label the cell membrane (cyan), and immunostained for tubulin (white). One z-section is shown. (C) Examples of cells in 2D (monolayers, top row) and 3D culture (spheroids, bottom row) of the indicated cell lines. Labeling as in B, but SiR-actin replaced WGA for all samples except RPE1 p53KD and OVSAHO monolayers. Images show maximum intensity projections, except MDA-MB-231 and OVSAHO spheroids, where one z-section is shown. (D) Percentage of mitotic cells in different phases (legend provided below the graph) for 4 cell lines in 2D or 3D culture, as indicated. (E) Examples of cells with unaligned or misaligned chromosomes (see schemes), labeling as in B. (F) Percentage of cells in late prometaphase and metaphase with different chromosome congression errors (see legend). The length of the scale bars is given in the images. In panels with multiple images, scale bars are equally long. In the graphs, the numbers of cells are displayed above each bar, with the statistical significance for the 2D versus 3D comparison shown at the top. Chi-square test was used. The number of independent replicates was 3–6 for 2D and 6–19 for 3D cultures.

Mitotic cells were observed throughout the spheroids, suggesting the absence of a necrotic core, likely due to the flattened spheroid shape. We estimated that mitotic cells comprised 2.97 ± 0.43% of cells in 2D monolayers in 8 independent experiments (129 mitotic cells out of 5289; data are presented as mean ± SEM over independent experiments) and 1.62 ± 0.37% in 3D spheroids in 6 independent experiments (32 mitotic cells out of 1912; Figure S1A). The difference was not significant (p = 0.052). Moreover, the fraction in 3D may be slightly underestimated because mitotic cells were harder to identify in crowded environments, suggesting that the true difference is likely even smaller. Culture conditions, however, influenced interphase cell morphology: cells grown in monolayers were mostly elongated or branched, whereas those in spheroids were predominantly round (Figure 1C)(Baker & Chen, 2012; Hakkinen et al., 2011; Knouse et al., 2018; McKenzie et al., 2018). These results indicate that, while 3D culture reshapes interphase cells, mitotic entry and overall cell cycle progression appear largely similar.

### Tumor cells in spheroids show a prometaphase delay and minor chromosome misalignment

Physical confinement can affect mitotic progression; for example, constrained growth of cells in spheroids embedded in agarose resulted in a prometaphase delay (Desmaison et al., 2013). To determine the effects of our spheroid models on the progression through mitosis, we quantified the distribution of mitotic phases and compared it to monolayer cultures. In RPE1 p53KD cells, there was no significant difference in the proportion of mitotic cells before and after anaphase onset between monolayers and spheroids (Figure 1D). In contrast, all 3 tumor cell spheroids had a significantly smaller fraction of mitotic cells that progressed to anaphase (Figure 1D). The largest difference was observed in MDA-MB-231 spheroids, where only 8% of mitotic cells were in phases after anaphase onset, compared to 23% in monolayers, suggesting difficulty in progressing into anaphase. Tumor spheroids, especially MDA-MB-231 and U2OS, showed a substantial increase in the fraction of prometaphase cells compared to monolayers, with approximately 60% of mitotic cells in this phase in spheroids (Figure 1D). In OVSAHO spheroids, the fraction of metaphase cells was increased. The accumulation of prometaphase cells in spheroids was not due to the nanomagnetic particles because the monolayer cultures were exposed to the same particles (see Methods). Overall, the elevated percentage of cells in prometaphase and metaphase in tumor spheroids suggests that these cells do not enter anaphase in a timely manner when compared to cells grown in monolayers, but instead activate the spindle assembly checkpoint (SAC) and prolong mitosis.

The observed prometaphase delay may be caused by inefficient chromosome alignment. To explore this possibility, we quantified the chromosomes that were not aligned at the metaphase plate, classifying them as unaligned chromosomes if they were found at or behind the spindle pole, or misaligned chromosomes if they were located between the pole and the metaphase plate (Figure 1E). In RPE1 p53KD and MDA-MB-231 spheroids, we found an increased fraction of mitotic cells that had unaligned or misaligned chromosomes, or both, in comparison with monolayers (Figure 1F). In U2OS and OVSAHO spheroids, the fraction of cells with chromosome alignment issues was similar in spheroids and monolayers (Figure 1F). Thus, inefficient chromosome alignment can partly explain the prometaphase delay observed in spheroids.

To test whether the delay in prometaphase affected chromosome segregation fidelity, we quantified chromosome segregation errors in anaphase and during cytokinesis. None of the cell lines showed an increase in anaphase or cytokinesis errors in spheroids compared to monolayers (Figure S1B–C). This suggests that during the prometaphase delay, the majority of chromosome alignment or attachment errors were corrected, and the cells proceeded to anaphase to divide mostly without errors.

### Mitotic cells in spheroids are rounder and slightly smaller than in monolayers

As interphase cells change their shape when grown in spheroids (Figure 1C), we hypothesized that the size and shape of mitotic cells may also be different from those in monolayers, which in turn may affect mitosis (Dix et al., 2018; O. M. Lancaster et al., 2013; Taubenberger et al., 2020). To quantify cell geometry, we measured the length and width of mitotic cells in the xy-plane, at the points of maximal and minimal extension, regardless of spindle orientation (Figure 2A). Both dimensions were significantly reduced in spheroids compared to monolayers in each cell line, except for width in RPE1 p53KD (Figure 2B, C). When pooled across all cell lines, cell length decreased from 27.53 ± 0.35 µm (n=571) in 2D to 19.98 ± 0.20 µm (n=464) in 3D (p=7.0 x 10^−73^), while cell width decreased from 20.87 ± 0.19 µm (n=571) in 2D to 16.34 ± 0.16 µm (n=464) in 3D (p=2.8 x 10^−66^). In spheroids, length and width were comparable within individual cells, consistent with a more rounded mitotic morphology, whereas in monolayers the greater separation between length and width indicated a more elongated shape in the xy-plane (Figure 2F; Figure S2A–D).

**Figure 2.**
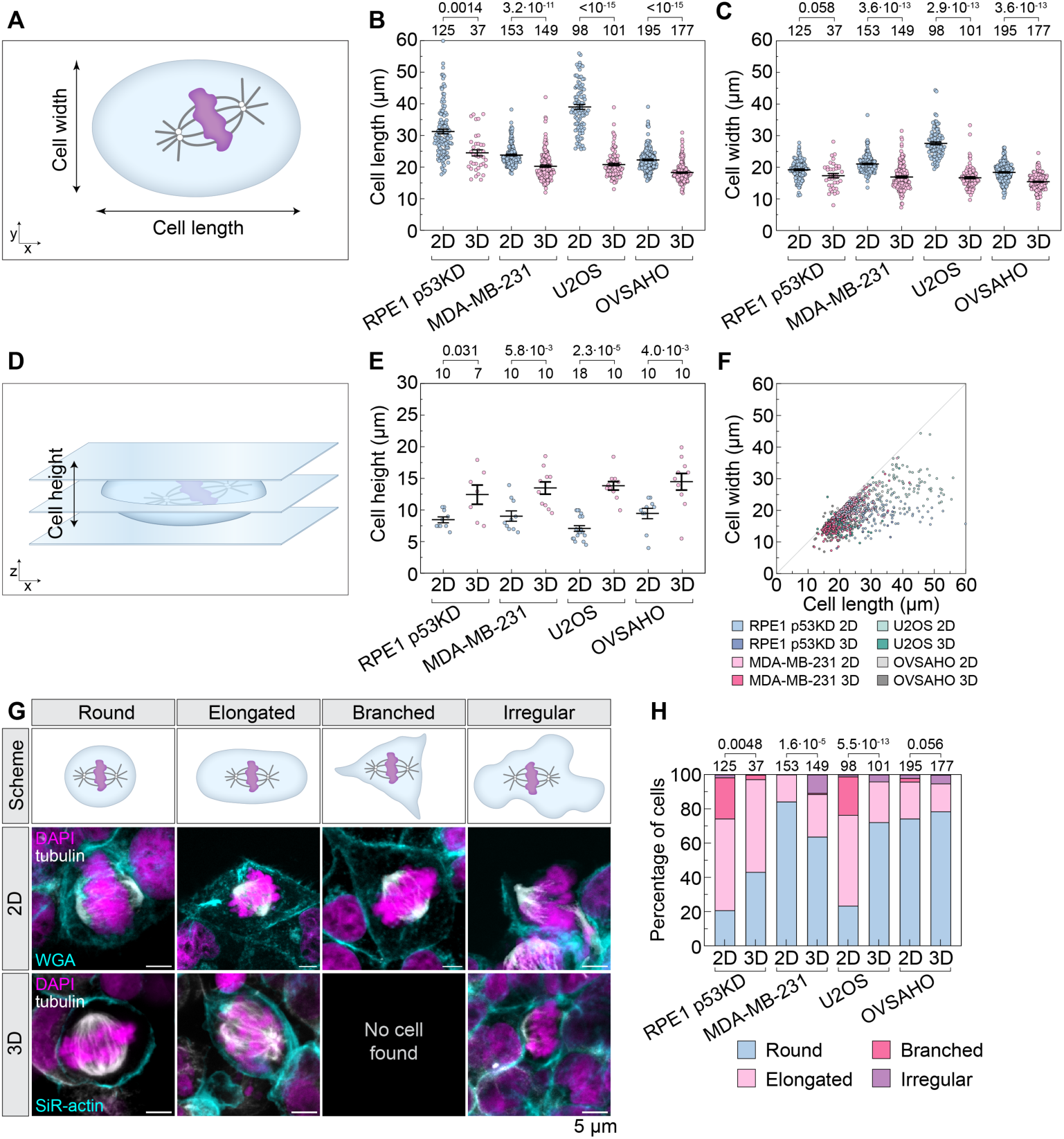
Mitotic cells in spheroids are smaller and more circular with shape that varies between cell lines. (A) Schematic representation of cell lengths and widths measured in mitotic cells. (B) Cell length and (C) cell width of cells in prometaphase, metaphase, or at the moment of anaphase onset, for 4 cell lines in 2D or 3D culture, as indicated. Mean and SEM values are shown, with each dot representing a single cell. (D) Schematic representation of cell height measured in mitotic cells. (E) Cell height of mitotic cells for 4 cell lines in 2D or 3D culture, as indicated. (F) Cell width as a function of cell length from the data points shown in B and C (see legend below for cell lines and culturing models). (G) Schematic representation of different cell shapes that appear during cell division, together with examples of different mitotic cell shapes in OVSAHO cell line monolayers and spheroids. DNA is labeled with DAPI (magenta), whereas cell membranes of monolayers and spheroids were stained with WGA and SiR-actin, respectively (both in cyan). Cells were also immunostained for tubulin (white). Images show the sum-intensity of all projections. The length of the scale bar is given below the images and has the same value for all images. (H) Percentage of different shapes in mitotic cells of different cell lines and culturing models (listed in legend below). In the graphs, Kruskal-Wallis test with post-hoc Dunn’s test was used for statistical analysis of B and C, the two-tailed Mann-Whitney U test was used in E to compare 2D and 3D cultures in the same cell lines, and Chi-square test was used for H.

To estimate the change in cell volume, we assessed cell height by counting the number of z-planes spanned by each cell in 2D and 3D cultures (Delarue et al., 2014). Cell height increased significantly from 2D to 3D in each cell line (Figure 2D, E). Across all cell lines together, it increased from 8.32 ± 0.33 µm (n=48) in 2D to 13.72 ± 0.55 µm (n=37) in 3D (p=7.7×10^−12^). Assuming a triaxial ellipsoid geometry, cell volume calculated from the measured dimensions decreased significantly from 2D to 3D only in U2OS cells (37%), whereas the other three cell lines showed no change, yielding a 17.26% overall reduction across all cell lines (p=0.032). These results indicate that cells transition from a flatter morphology in 2D to a more spherical morphology in 3D, accompanied by a decrease in volume only in U2OS cells. To quantify the specific changes in cell shape in the xy-plane, cells in prometaphase, metaphase, and at anaphase onset were classified as round, elongated, branched, or irregular (Figure 2G, H). U2OS cells showed the largest shift from elongated shapes in 2D to round in 3D, RPE1 p53KD cells also became rounder, MDA-MB-231 developed irregular shapes, and OVSAHO remained unchanged (Figure 2H). Collectively, these data show that mitotic cells are rounder in spheroids than in monolayers, and the specific morphologies vary by cell line.

### Mitotic spindles in spheroids are smaller and prone to multipolarity, misalignment, and displacement

The altered cell size and shape in spheroids may affect spindle size and shape. To investigate spindle properties in spheroids, we first measured spindle length and width in metaphase cells (Figure 3A–C). We found a significant reduction in both spindle dimensions across all cell lines cultured as spheroids compared to monolayers, except for spindle width in RPE1 p53KD, where no significant difference was detected (Figure 3B-D, Figure S3A–D). The most pronounced reduction in spindle length was found in U2OS cells (Figure S3C), which also showed the greatest reduction in cell length (Figure 3E), and were the only cell line with a significant decrease in cell volume. These results indicate that spindle length adapts to the reduced cell size in spheroids.

**Figure 3.**
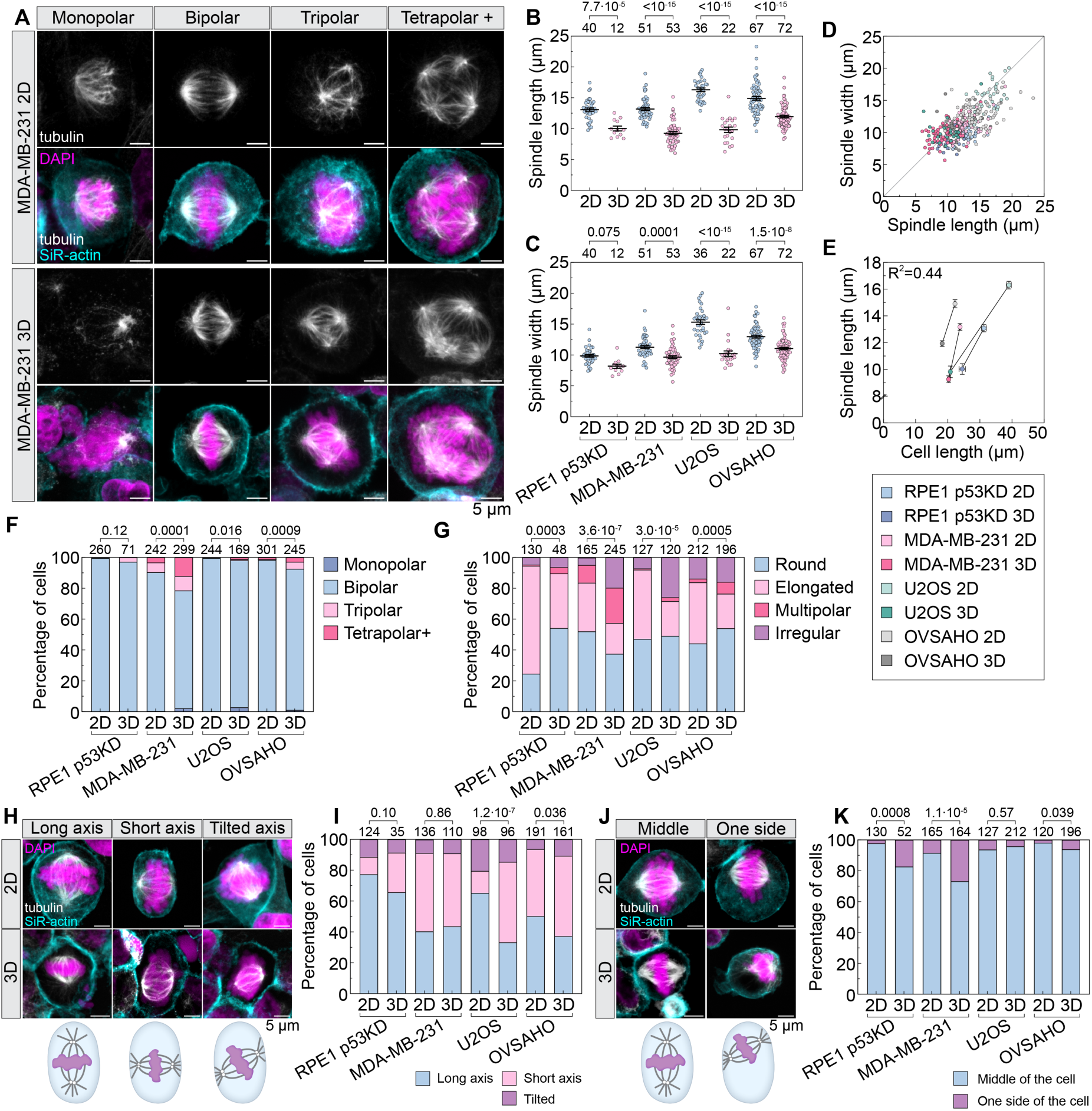
Mitotic spindle in spheroids has higher polarity and exhibits different size, position, and orientation compared to monolayers. (A) Examples of different spindle polarities in cells that compose monolayers or spheroids, stained with DAPI to label DNA (magenta), SiR-actin to label actin cytoskeleton (cyan), and immunostained for tubulin (white). Images show the sum of all intensity projections. (B) Spindle length and (C) spindle width of bipolar metaphase cells, for 4 cell lines in 2D or 3D culture, as indicated. Graphical representations show mean and SEM values, with each dot representing a single cell. (D) Association of spindle length with spindle width and (E) cell length (see Figure 2B). The different combinations of cell lines and cell culture models are shown in the legend below. (F) Percentage of cells in late prometaphase and metaphase with spindle polarities indicated on the legend. Monolayers and spheroids in 4 different cell lines were compared. (G) Percentage of a particular spindle shape (see legend) in monolayers and spheroids, compared in 4 different cell lines. (H) Examples of different spindle orientations in MDA-MB-231 monolayers and spheroids (labeling as in A), with corresponding schematic representations illustrated below. Images show sum of all intesity projections. (I) Percentage of cells with a particular spindle orientation, compared between monolayers and spheroids in 4 different cell lines. (J) Examples of different spindle positioning in MDA-MB-231 monolayers and spheroids (labeling as in A and H), with corresponding schematic representations illustrated below. Images show the sum of all Z-slices. (K) Percentage of cells with a particular spindle orientation, compared between monolayers and spheroids in 4 different cell lines. The scale bar lengths are given and have the same value for all images in panels. Kruskal-Wallis test with post-hoc Dunn’s test was performed for B and C. Pearson’s correlation coefficient was used in E. Fisher’s exact test when compared groups had two outcomes, and Chi-square test if there were more than two outcomes in F. Chi-square test was performed in G and I, and Fisher’s exact test in K.

The fraction of multipolar spindles increased in tumor cell spheroids compared to monolayers, with the most prominent increase in MDA-MB-231 cells (Figure 3A,F). A small fraction of monopolar spindles was also detected in tumor spheroids (Figure 3A,F). An extended analysis of spindle shapes, where spindles were categorized as round, elongated, multipolar, or irregular, revealed changes in spindle shape in all cell lines grown in spheroids compared to monolayers (Figure 3G, Figure S3E). RPE1 p53KD spheroids showed an increase in round spindles, MDA-MB-231 in multipolar and irregular ones, U2OS in irregular ones, and OVHASO in round and multipolar spindles (Figure 3G). In MDA-MB-231 spheroids, multipolar and irregular spindles were more frequently found in irregularly shaped cells than in round or elongated ones (Figure S3F), indicating that specific changes in spindle morphology reflect particular changes in cell shape.

To test whether spindles in spheroids follow Hertwig’s rule (Hertwig & Hertwig, 1884), which states that spindles align along the long axis of the cell, we used bipolar spindles and quantified the fraction of them that were aligned with the cell’s long axis, short axis, or tilted (Figure 3H). RPE1 p53KD cells followed Hertwig’s rule in spheroids as well as in monolayers, whereas most of the MDA-MB-231 cells did not follow this rule in both spheroids and monolayers (Figure 3I). Spindles in U2OS cells were aligned according to Hertwig’s rule in monolayers, but shifted towards the alignment along the short cell axis in spheroids, and OVSAHO cells showed a similar but smaller shift (Figure 3I). These results indicate that spindle orientation in spheroids is cell line-specific and can deviate from classical geometric rules in the 3D environment.

Spindles were predominantly positioned centrally within the cell in all lines in spheroids as well as in monolayers, but in a fraction of cells, the spindle was displaced off-center, close to the cell membrane (Figure 3J). The fraction of off-centered spindles was larger in spheroids than in monolayers in RPE1 p53KD and MDA-MB-231 cells, and there was a minor increase in OVSAHO cells (Figure 3K). Overall, our analysis of spindle geometry and position shows that spindles in spheroids are smaller, more frequently multipolar and irregularly shaped, as well as misaligned and displaced, in comparison with spindles in monolayers.

### Mitotic proteins are downregulated in cancer spheroids

To assess the changes in the global proteome between 2D and 3D cultures, we conducted quantitative proteomic analysis on whole-cell lysates from RPE1 p53KD, MDA-MB-231, U2OS, and OVSAHO cells grown as 2D monolayers or 3D spheroids. Principal component analysis (PCA) showed distinct clustering of samples according to culture condition, confirming overall proteomic differences between 2D and 3D cultures (Figure S4A). Across most conditions, approximately 6,000 protein groups were identified per sample, whereas RPE1 p53KD 3D samples displayed a lower number of proteins (∼4,000) (Figure S4B). The total summed protein intensity for this sample was higher (1 × 10^9) compared to the other conditions (≈7 × 10^8) (Figure S4C).

Quantitative analysis of abundance ratios (3D/2D) showed a broad downregulation of proteins annotated with mitotic Gene Ontology (GO) terms in 3D cultures (Figure 4A). Downregulation of mitotic proteins was particularly evident in U2OS and MDA-MB-231 cells, whereas RPE1 p53KD cells showed a weaker overall reduction. Proteins upregulated in 2D cultures were primarily associated with nuclear transport and RNA processing, whereas those upregulated in 3D cultures were enriched for mitochondrial and metabolic functions (FigureS4D–I). There were no major changes in Ki-67, a marker for cell proliferation that is primarily expressed in active cell cycle phases (S, G2, M) but is progressively degraded in G1 and absent in G0 phase (Miller et al., 2018; Sobecki et al., 2017; Uxa et al., 2021) (Figure S4J). A schematic representation of cell-cycle proteins illustrates an overall reduction of mitotic regulators in 3D versus 2D, with each protein color-coded by 3D/2D abundance ratio and quadrants representing each cell line (Figure 4B). Among key cell-cycle regulators, CCNA2 (Cyclin A2), which coordinates DNA replication and entry into mitosis (Bouhamida et al., 2025; Silva Cascales et al., 2021), was significantly reduced in 3D cultures of U2OS and RPE1 p53KD (Figure 4C). A significant reduction across all cell lines was observed for CCNB1 (Cyclin B1), a key regulatory protein that triggers entry into mitosis (Jin et al., 1998; Maryu & Yang, 2022; Pfaff & King, 2013) (Figure 4C). Spindle assembly checkpoint (SAC) proteins, such as BUB1B, were generally downregulated in spheroids, except MAD1L1, which was upregulated in RPE1 cells (Figure 4C, Figure S4J). Components of the anaphase-promoting complex/cyclosome (APC/C), including ANAPC5 and CDC20, were downregulated in 3D cultures across multiple cell lines (Figure 4C, Figure S4J).

**Figure 4.**
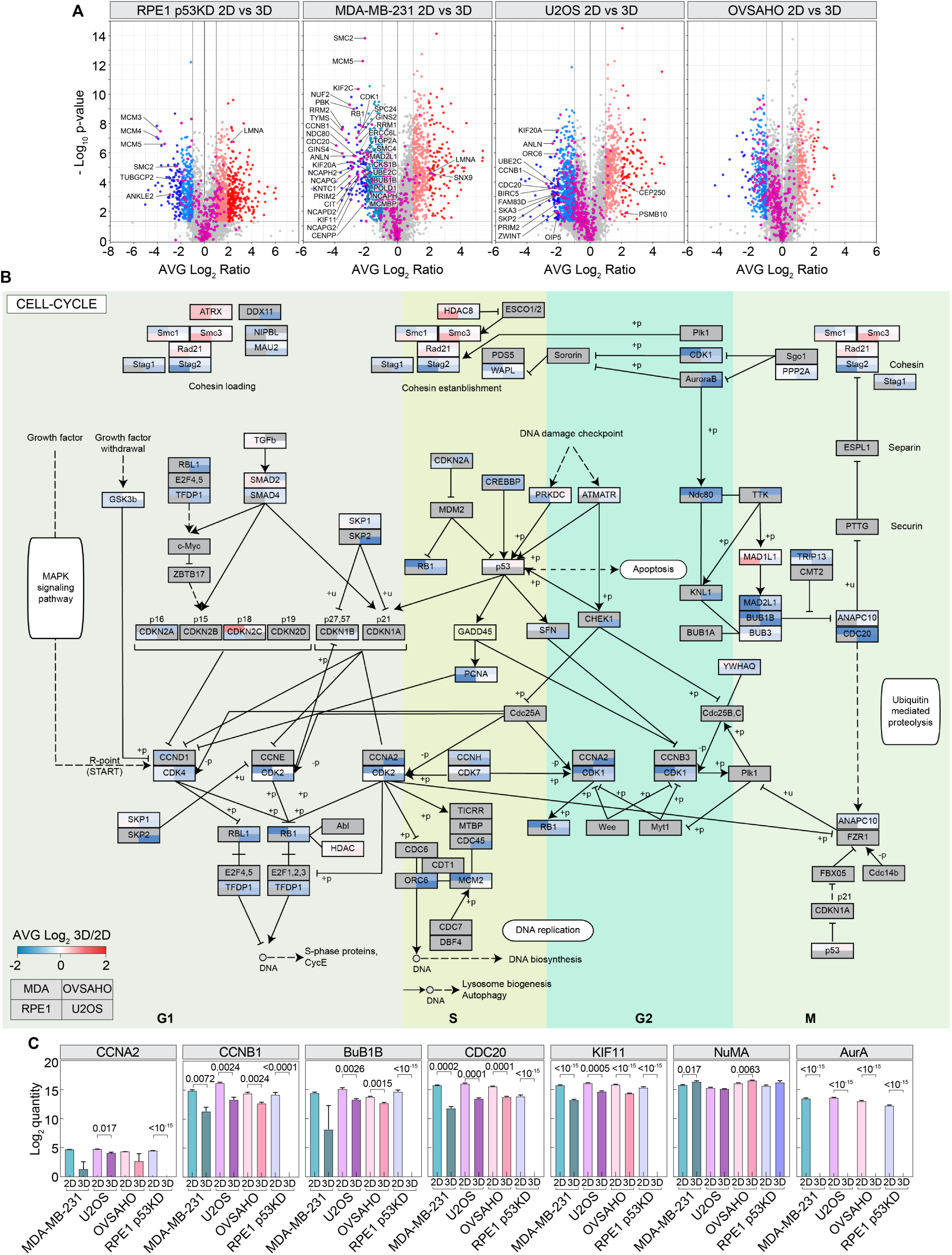
Spheroids display altered expression of mitotic proteins. (A) Volcano plot of log2-transformed ratios for proteins quantified in the 3D culture samples versus in the 2D culture samples of RPE1 p53KD, MDA-MB-231, OVSAHO, and U2OS cell lines. Significant proteins are defined as p < 0.05, with class A proteins showing log₂ fold change >2 and class B proteins >1. Class A upregulated proteins are color-coded in red, class B in salmon, class A downregulated proteins are color-coded in dark blue, and class B in light blue. Proteins marked in pink are those annotated as mitotic according to GO terms, with those that are also class A listed with their gene name. (B) Schematic illustration of the ratio between 3D culture and 2D culture of proteins involved in the cell cycle according to KEGG Pathway Database. Proteins are color-coded with a gradient from blue (more abundant in 2D) to red (more abundant in 3D). Each circle represents one protein and is divided into four parts, each representing one of the cell lines. Proteins with no quantifiable ratios are shown in dark grey. (C) Log2 quantity of different proteins in 2D and 3D cultures across all cell lines. The mean of three technical replicates (n = 3) ± SEM is shown. A t-test was used to compare 2D versus 3D (p-values are indicated in the plots).

Among the proteins that regulate spindle size and position, we found that KIF11, a kinesin essential for spindle bipolarity and length-regulation (Vukušić et al., 2021; Silva et al., 2025), was consistently downregulated in 3D cultures of all four cell lines (Figure 4C). KIF4A, another kinesin involved in chromosome condensation and anaphase spindle elongation (Vukušić et al., 2021), followed a similar pattern, though the decrease was significant only in OVSAHO cells (Figure S4J). NUMA, one of the proteins that link the spindle to the cell cortex (Cho et al., 2025; Seldin et al., 2013), was upregulated in 3D in all cell lines except U2OS (Figure 4C). Together, these findings suggest that cells grown in 3D downregulate a network of mitotic regulators and shift their proteomic profile toward mitochondrial and metabolic pathways, which is consistent with altered mitotic behavior and spindle morphology in spheroids.

## Discussion

In this work, we demonstrate that cell growth in a 3D culture remodels mitosis at both structural and proteomic levels, revealing shared and cell line-specific adaptations to spheroid culture (Figure 5). Across four epithelial lines spanning non-transformed and tumor origins, spheroid culture produced rounder mitotic cells with shorter spindles, increased multipolar and irregular spindle morphologies, and led to more frequent spindle misalignment and off-center positioning. We found accumulation of prometaphase and metaphase cells, indicative of a mitotic delay due to the activation of the spindle assembly checkpoint. The phenotypes we observed are consistent with prior evidence that physical confinement prolong prometaphase and perturb spindle positioning and integrity (Charnley et al., 2013; Desmaison et al., 2013, 2018; Molla et al., 2017). Notably, despite prolonged prometaphase and elevated alignment defects in some lines, chromosome segregation errors in anaphase remained low, implying that extended prometaphase allows for the correction of most early attachment errors. Together, our findings argue that spatial architecture and mechanical context are important determinants of mitotic progression in tumors, altering spindle geometry, orientation, and timing in a 3D setting, without necessarily increasing final segregation errors when the checkpoint is functional and cells have sufficient time for correction.

**Figure 5.**
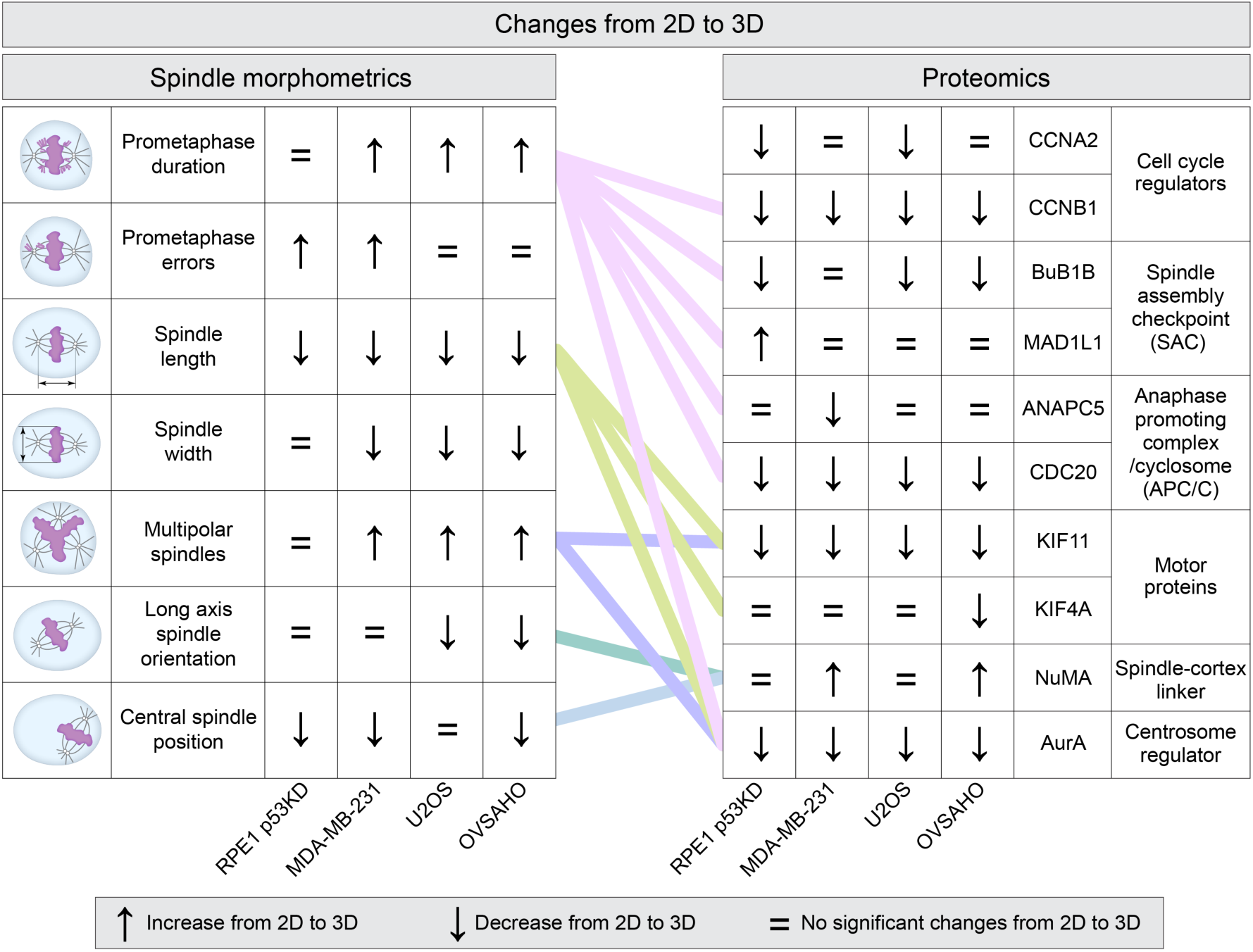
Summary of mitotic phenotypes and proteomic changes in 3D cultures. Mitotic phenotypes observed in 3D spheroids relative to 2D monolayers (left) and corresponding changes in protein abundance identified by quantitative proteomics (right). Connecting lines represent inferred relationships between protein-level changes and observed mitotic phenotypes.

The proliferation rate of cells in spheroids was not changed compared to monolayers, as the fraction of mitotic cells determined by microscopy and Ki-67 levels determined by quantitative proteomics were largely unchanged between 2D and 3D cultures. This indicates that cells in spheroids remain actively cycling. Quantitative proteomics revealed a broad downregulation of mitotic regulators in 3D cultures, together with enrichment of metabolic and mitochondrial pathways, an overall shift that was previously observed in comparative 2D–3D proteomics (Kumar et al., 2008). We observed a reduction of core cyclins in 3D cultures, with Cyclin A2 downregulated in U2OS and RPE1 p53KD cells and Cyclin B1 consistently reduced across all cell lines. Together with decreased levels of spindle assembly checkpoint and APC/C components, these changes indicate a global rewiring of cell-cycle regulation in 3D.

The decreased abundance of APC/C components in 3D is expected to lead to a reduced efficiency of metaphase-anaphase transition (Wild et al., 2016), which is consistent with the observed prometaphase delay, increasing opportunities for the correction of transient spindle multipolarity and chromosome congression defects. The observed prometaphase delay suggests that, even though spindle assembly checkpoint proteins were broadly downregulated in 3D cultures, the extent of this reduction appears to remain within the functional range of the checkpoint.

Changes in spindle-associated motors and cortical regulators further link molecular regulation to altered spindle geometry. Downregulation of the mitotic kinesins KIF11 and KIF4A, which control spindle length in metaphase and elongation in anaphase (Vukušić et al., 2021), likely contributes to reduced spindle size and increased spindle abnormalities. Upregulation of NUMA, one of the proteins that link the spindle to the cell cortex (Sun et al., 2021), may compensate for altered force balance and spindle positioning in rounder cells. Overall, our findings indicate that the growth environment shapes mitosis through a combination of mechanical cues that directly influence spindle architecture and intrinsic proteomic changes associated with spheroid growth, which may indirectly modulate mitotic progression.

The heterogeneity across cell lines highlights how genomic background and baseline architecture modulate 3D responses. Among the tested cell lines, the most mesenchymal-like line, MDA-MB-231, exhibited the largest changes in 3D versus 2D, including a high prometaphase fraction, spindle multipolarity, and misorientation. This inter-line variation is consistent with reports that confinement-induced mitotic delays and spindle orientation defects depend on cell shape and the capacity for mitotic rounding (Charnley et al., 2013), which differ across tumor types and matrix conditions.

Our integrated 2D–3D cross-line analysis addresses a key gap in the field by pairing detailed mitotic morphometrics with unbiased proteomics in matched samples, complementing earlier studies that profiled global proteome differences without simultaneous mitotic phenotyping (Kumar et al., 2008) or those that focused solely on mechanics and imaging (Desmaison et al., 2018). Based on our results, future studies may include targeted perturbations, such as partial restoration of KIF11/KIF4A levels or APC/C activity in spheroids, to determine whether mitotic delay, multipolarity, and misalignment can be rescued under 3D culture. Similarly, controlled modulation of 3D mechanics, by varying spheroid shape or matrix stiffness, could dissect how mechanical inputs and proteomic remodeling jointly set the thresholds for spindle integrity and checkpoint satisfaction in different culture contexts. Moving forward, integrating single-cell proteomics (Ye et al., 2025) with live imaging in 3D cultures and controlled mechanical perturbations should resolve how cell-to-cell variability in rounding, spindle-cortex coupling, and mitotic proteome dictates mitotic progression and fidelity.

By establishing a framework that connects proteome state to mitotic architecture in 3D, this study provides a foundation for identifying mitosis-specific vulnerabilities that are amplified in 3D environments but quiet in 2D monolayers. These insights have implications for the development of new therapies and tumor model selection. Incorporating 3D assays and readouts of mitotic architecture and timing, alongside proteomic signatures of mitotic regulators, may lead to more efficient and context-aware strategies to target mitosis in cancer.

## Methods

### Cell culture

Human hTERT RPE1 cell line with a p53 knockdown (KD) stably expressing H2B-Dendra2 (referred to as RPE1 p53KD) (Soto et al., 2018), a gift from Rene Medema (Netherlands Cancer Institute, Amsterdam, Netherlands), unlabeled human breast cancer MDA-MB-231 cell line, a gift from Dragomira Majhen (Ruđer Bošković Institute, Zagreb, Croatia), unlabeled human osteosarcoma U2OS cell line, a gift from Marin Barišić (Danish Cancer Society Research Center, Copenhagen, Denmark) and unlabeled human high-grade serous ovarian cancer OVSAHO cell line (JCRB1046 Cell Bank, Tebubio, Le Perray-en-Yvelines, France) were used. All cells were maintained in Dulbecco’s Modified Eagle Medium (DMEM) medium (Gibco, MT, USA), except OVSAHO, which were kept in RPMI-1640 medium (Sigma-Aldrich, MO, USA). Cell culture medium was supplemented with 10% heat-inactivated fetal bovine serum (FBS, Sigma-Aldrich, MO, USA) and penicillin (100 IU/mL)/streptomycin (100 mg/mL) solution (Lonza, Basel, Switzerland). Cells were grown in a Galaxy 170S humidified incubator (Eppendorf, Hamburg, Germany) at 37°C with a 5% CO_2_ atmosphere. Cells were regularly tested for mycoplasma contamination with DAPI staining.

### Spheroid generation using magnetic cell levitation

Spheroids were made by the magnetic cell levitation method (Souza et al., 2010) with 6 Well Bio-Assembler^TM^ Kit (Greiner Bio-One GmbH, Kremsmünster, Austria). To achieve the optimal confluency of approximately 70% for treatment with a nanoparticle solution, 250000 cells were seeded as monolayers one day prior to the addition of 1 µL/cm^2^ of the nanoparticle solution. After 18 hours of incubation, the cell suspension was transferred into a cell-repellent 6-well plate with a magnetic lid. Spheroids were incubated for 5–7 days before immunostaining or sample preparation for proteomics. Monolayers were also incubated with nanoparticles, and a magnetic lid was placed over the cells for 4 hours, but not longer to avoid the formation of spheroids.

### Immunofluorescence

Samples were fixed with 4% paraformaldehyde for 20 min at room temperature. Next, a blocking buffer containing 2% NGS and 0.5% Triton-X-100 in PBS was added and incubated for 1 hour at room temperature. Primary antibody rat anti-tubulin (MA1-80017, Invitrogen) was diluted in a 1:300 ratio in blocking buffer and incubated overnight at 4°C. On the following day, the secondary antibody, donkey anti-rat Alexa Fluor 488 (ab150153, Abcam) or donkey anti-rat Alexa Fluor 647 (ab150155, Abcam), diluted in a 1:500 ratio in the blocking buffer, was incubated for 1 hour at room temperature, in the dark. Samples were stained with 1 µg/mL DAPI to label the DNA and 4 µM SiR-actin (Spirochrome, Stein am Rhein, Switzerland) or 5 µg/mL Wheat Germ Agglutinin (WGA) Alexa Fluor^TM^ 594 Conjugate (Invitrogen, Waltham, MA, SAD) to label the cell membrane/cortex, and incubated for 20 minutes. The samples were washed with PBS for 10 min between each step, and finally mounted with antifade aqueous embedding media (Abberior, Göttingen, Germany) and covered with a coverslip.

### Microscopy

Confocal imaging was performed on an Airyscan Zeiss LSM800 confocal laser scanning microscope with the 63x/1.4 Oil DICII objective (Carl Zeiss, Germany) and LSM 800 camera. Laser lines of 405 nm, 488 nm, 561 nm, and 640 nm were used to excite DAPI, Alexa Fluor 488, 568, 594, and 647, respectively. The number of z-slices was adjusted individually for each cell during spheroid imaging, to capture the whole cell, and for monolayers, 31 z-slices were imaged per cell. Only 0.3, 0.5 or 1.0 µm z-step sizes were used in experiments. Images were acquired in ZEN Blue 3.5 software.

### Image analysis

Images were analyzed in Fiji/ImageJ (National Institutes of Health, Bethesda, MD, USA) and ZEISS ZEN 3.13 software (Carl Zeiss, Germany). The percentage of mitotic cells was obtained by counting all interphase and mitotic cells in the chosen z-slice. Only one z-plane was analyzed in 2D, whereas 3D data were obtained from three distinct z-planes separated by 30 planes to minimize the possibility of counting the same cell multiple times. When insufficient number of planes was available, only one plane was analyzed.

Cells were categorized according to their mitotic phase: prophase, prometaphase, metaphase, anaphase, telophase, or cytokinesis. Prophase was identified by the DNA that had begun to condense into chromosomes, while the nuclear shape was still visible. Early prometaphase was defined as the phase in which all chromosomes were fully condensed, and the mitotic spindle had begun to form, with clearly visible spindle poles and the spindle body. Cells in early prometaphase were excluded from the analyses of cell and spindle shape and size, because the spindle was not yet fully formed, and the cell could still round up later in prometaphase. Late prometaphase was defined as the phase in which the mitotic spindle was fully formed, and all chromosomes were located on the spindle, but not yet aligned at the metaphase plate. Metaphase was defined as the phase in which all, or nearly all, chromosomes were aligned at the equatorial plane. Anaphase was defined as the phase in which chromosomes had begun to separate, but the cell had not yet started to divide into two daughter cells, i.e., the actomyosin ring had not yet formed. Telophase was defined as the phase in which chromosomes were well separated and still largely condensed, but the two daughter cells had begun to separate with the formation of an actomyosin ring. Cytokinesis was defined as the phase in which chromosomes had decondensed, daughter cells had formed, but remnants of the actomyosin ring were still visible.

Mitotic errors were defined as follows. If one or several chromosomes were located at the spindle pole during late prometaphase or metaphase, they were classified as unaligned chromosomes. If one or several chromosomes were located between the spindle pole and the metaphase plate and were not associated with the rest of the chromosomes, they were classified as misaligned chromosomes. If one or more chromosomes lagged between the two segregating chromosome masses during anaphase and were not connected to the others, they were classified as lagging chromosomes. If such a chromosome persisted through telophase and cytokinesis without being incorporated into the nucleus of the daughter cell, it was defined as a micronucleus. If one or more chromosomes remained connected to both segregating chromosome masses during anaphase, they were classified as chromosome bridges. Such chromosome bridges were also defined in cytokinesis, where they connected the two segregating chromosome masses through the actomyosin ring.

Cell length and width were measured only when the whole cell was in the field of view. Cell height was calculated by multiplying the difference between cell’s top and bottom z-planes with the z-step size. Effective cell volumes were calculated by multiplying their length, width, and height. Spindle length and width were measured if the spindle was horizontal relative to the dish surface.

### Sample preparation for mass spectrometry

Cell pellets were lysed utilizing a urea lysis buffer (6 M urea, 2 M thiourea, 60 mM Tris pH 8.0) and maintained on ice for 20 minutes. Subsequently, DNA and RNA were eliminated using benzonase (1 U/ml, Merck Millipore) for 10 minutes at room temperature. The samples underwent centrifugation for 10 minutes at 13,000 rpm and 4°C. Protein quantification was executed employing Bradford reagent, utilizing a standard curve with known concentrations of bovine serum albumin (BSA). The absorbance was assessed at 595 nm. Proteins were subjected to reduction with 10 mM dithiothreitol (DTT) for one hour, followed by alkylation with 55 mM iodoacetamide for one hour, and subsequent digestion with Lys-C (Lysyl Endopeptidase, Wako Chemicals) for three hours at room temperature. Following this, four volumes of 10 mM ammonium bicarbonate were added, and proteins were digested with trypsin (Promega Corporation) overnight. Digestion was halted by the addition of 1% trifluoroacetic acid (TFA).

### Liquid chromatography - mass spectrometry measurement

Samples were analyzed using an Exploris 480 mass spectrometer (Thermo Fisher Scientific) coupled online with an Easy-nLC 1200 UHPLC system (Thermo Fisher Scientific). Equal amounts of peptides were loaded onto a 20-cm-long, 75-µm inner-diameter PicoTip fused-silica analytical column (New Objective) packed in-house with ReproSil-Pur C18-AQ 1.9-µm silica beads (Dr. Maisch GmbH). The column was maintained at 40 °C. Peptides were loaded at a flow rate of 1 µL/min and separated over a 60-minute linear gradient from 4% to 95% of solvents A (0.1% formic acid) and B (80% acetonitrile in 0.1% formic acid) at a steady flow rate of 300 nL/min. Electrospray ionization was conducted at 275 °C in positive ion mode.

Data were acquired in data-independent acquisition (DIA) mode. Full MS1 scans were acquired over an m/z range of 400–1000 at a resolution of 120,000 with the automatic gain control (AGC) set to “custom” (normalized value 300%, absolute target 3.0 × 10⁶). Sixty contiguous DIA windows of 10 m/z were used to cover the precursor range, with the window placement optimization on. MS2 scans were acquired using HCD collision energy at 27%, over an m/z range of 145–1450, at a resolution of 15,000, with AGC set to “custom” (normalized value 800%, absolute target 8.0 × 10⁵).

### Mass spectrometry data processing

Raw data were processed using Spectronaut software (version 19.1.240806.62635; Biognosys AG) with standard parameters as recommended by the manufacturer. Spectra were searched against a Uniprot Homo sapiens database (104 556 entries, downloaded on 2024/01/30). Downstream analysis on the Protein report was performed in Excel and Perseus software (Tyanova et al., 2016). Proteins were functionally annotated with Gene Ontology (GO) Biological Processes, GO Cellular Compartment, GO Molecular Functions, and Kyoto Encyclopedia of Genes and Genomes. Significance was defined as p <0.05, and class A proteins as those showing log₂ fold change >2 and class B proteins >1. The Fisher’s exact test (FDR ≤ 0.5) was used to assess the over-represented categories in Figure S4F,I; it was performed with the PANTHER Classification System software available online at https://www.pantherdb.org. For generating Venn diagrams, the online tools https://www.stefanjol.nl/venny and https://bioinfogp.cnb.csic.es/tools/venny/index.html were used. Box plot analysis of label incorporation was conducted with the online tool: http://shiny.chemgrid.org/boxplotr/. Additional graphical visualization was performed using the R environment (version 4.1.1) and GraphPad (version 8.0.1).

### Statistical analysis

Results for cell biology analyses were obtained from the following number of independent experiments for monolayers: RPE1 p53KD – 3, MDA-MB-231 – 5, U2OS – 6, and OVSAHO – 3; and for spheroids: RPE1 p53KD – 19, MDA-MB-231 – 15, U2OS – 6, and OVSAHO – 6. Quantification and statistical analysis were performed in GraphPad Prism (GraphPad Software, Boston, MA, USA). No statistical methods were used to predetermine the sample size. Data are given as mean ± SEM (standard error of the mean), unless stated otherwise. The data were tested for normal distribution using the Shapiro-Wilk normality test. The data were normally distributed for spindle length, spindle width, and cell width, whereas cell length did not have a normal distribution. When the data were normally distributed, one-way ANOVA followed by Tukey’s multiple comparison test was used for comparisons among multiple groups. If the data were not normally distributed, multiple groups were tested with the Kruskal-Wallis test followed by Dunn’s multiple comparison test. Proportions among the groups were statistically compared with the Chi-Square test, or Fisher’s exact test when the proportions of two outcomes were compared. A p-value < 0.05 was considered statistically significant. Figures and schemes were assembled in Adobe Illustrator CS5 and CC (Adobe Systems, Mountain View, CA, USA).

## Acknowledgements

We thank Rene Medema, Dragomira Majhen, and Marin Barišić for providing cell lines; Ivana Šarić for illustrations and figure assembly; Petar Škrobo for assistance with final manuscript preparation; and members of the Tolić lab for their constructive feedback. The work in the Tolić lab was funded by the European Research Council (ERC-SyG 855158), the Croatian Science Foundation (HRZZ IP-2024-05-5336), the Swiss-Croatian Bilateral Project (IPCH-2022-10-9344); and projects co-financed by the Croatian Government and the European Union through the European Regional Development Fund—the Competitiveness and Cohesion Operational Programme IPSted (KK.01.1.1.04.0057) and QuantiXLie Center of Excellence (KK.01.1.1.01.0004). The work of doctoral students An.P. and Ad.P. has been supported in part by the “Young researchers’ career development project – training of doctoral students” of the Croatian Science Foundation. C.C.-R. and B.M. were supported by the German Research Foundation (DFG) through GRK2364.

## Author contributions

I.M.T. and B.M. designed the research and supervised the cell biology and proteomics work, which was carried out by An.P. and C.C.-R., respectively. Ad.P. performed additional analysis of microscopy data. I.M.T. and An.P. wrote the paper, with help from Ad.P., C.C.-R., and B.M.

## Data availability

Relevant data supporting the findings of this study are provided within the article and its supplementary files. Proteomics raw data will be uploaded to the PRIDE database (Perez-Riverol et al., 2025) upon acceptance of the manuscript.

