## Supplementary figures for "Spheroid culture remodels mitosis and the proteome in tumor cells"

### Supplementary information

Ana Petelinec[1*], Claudia Cavarischia-Rega[2*], Adrian Perhat[1], Boris Maček[2], Iva M. Tolić[1#]

[1]Division of Molecular Biology, Ruđer Bošković Institute, Zagreb, Croatia
[2]Quantitative Proteomics, Department of Biology, Interfaculty Institute of Cell Biology, University of Tübingen, Germany
[*]Equal contribution

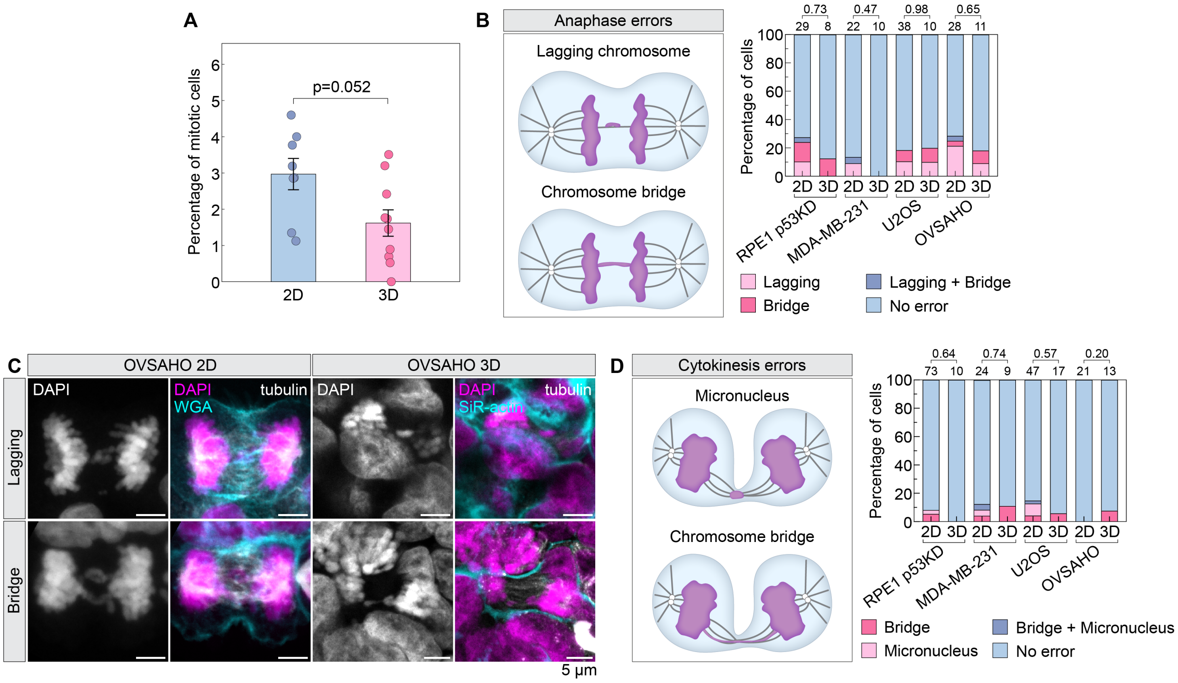


**Supplementary figure 1. Percentage of mitotic cells together with anaphase and cytokinesis errors does not differ between monolayers and spheroids**

(A) Percentages of mitotic cells in 2D and 3D cultures. Each dot represents one replicate, which was measured in one z-plane. Only MDA-MB-231 and OVSAHO cells were included in the analysis. MDA-MB-231 cells used for the analysis of 2D samples were expressing H2B-GFP, unlike the wild-type cells used for the 3D culture. OVSAHO cells used for the analysis of 2D samples were labeled with 10 nM SPY 555 dye, together with 1 µM verapamil, unlike the wild-type cells used for the 3D culture. Cells in spheroids were counted in one z-plane or, in cases where multiple z-planes were imaged, in three z-planes spaced far enough to avoid counting the same cells twice. Therefore, multiple technical replicates were included for one biological replicate, where possible. Only one z-plane was examined in 2D cultures.

(B) Schematic representation of two different anaphase error types, together with their proportions in four different cell lines.

(C) Examples of different anaphase error types in OVSAHO monolayers and spheroids. DNA is labeled with DAPI (white and magenta), with immunostained tubulin (white), whereas WGA and SiR-actin are used to label cell edges is monolayers and spheroids respectively (cyan). Images show sum-intesity of all projections.

(D) Schematic representation of two different cytokinesis error types, together with their proportions in four different cell lines. The length of the scale bars is given in the images. In (A)Two-sample Z-test for Proportions was performed to determine the statistical significance and the corresponding p-value.

In (B) and (D), Chi-square test was performed to determine the statistical differences and corresponding p-values.

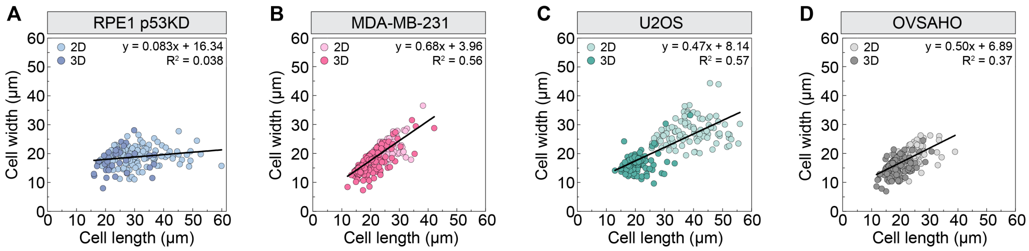


**Supplementary figure 2. Cells that compose spheroids are smaller with more uniform cell length and width than in monolayers**

Cell width as a function of cell length in monolayers and spheroids stated for RPE1 p53KD (A), MDA-MB-231 (B), U2OS (C), and OVSAHO (D) cell lines. Each dot represents one cell.

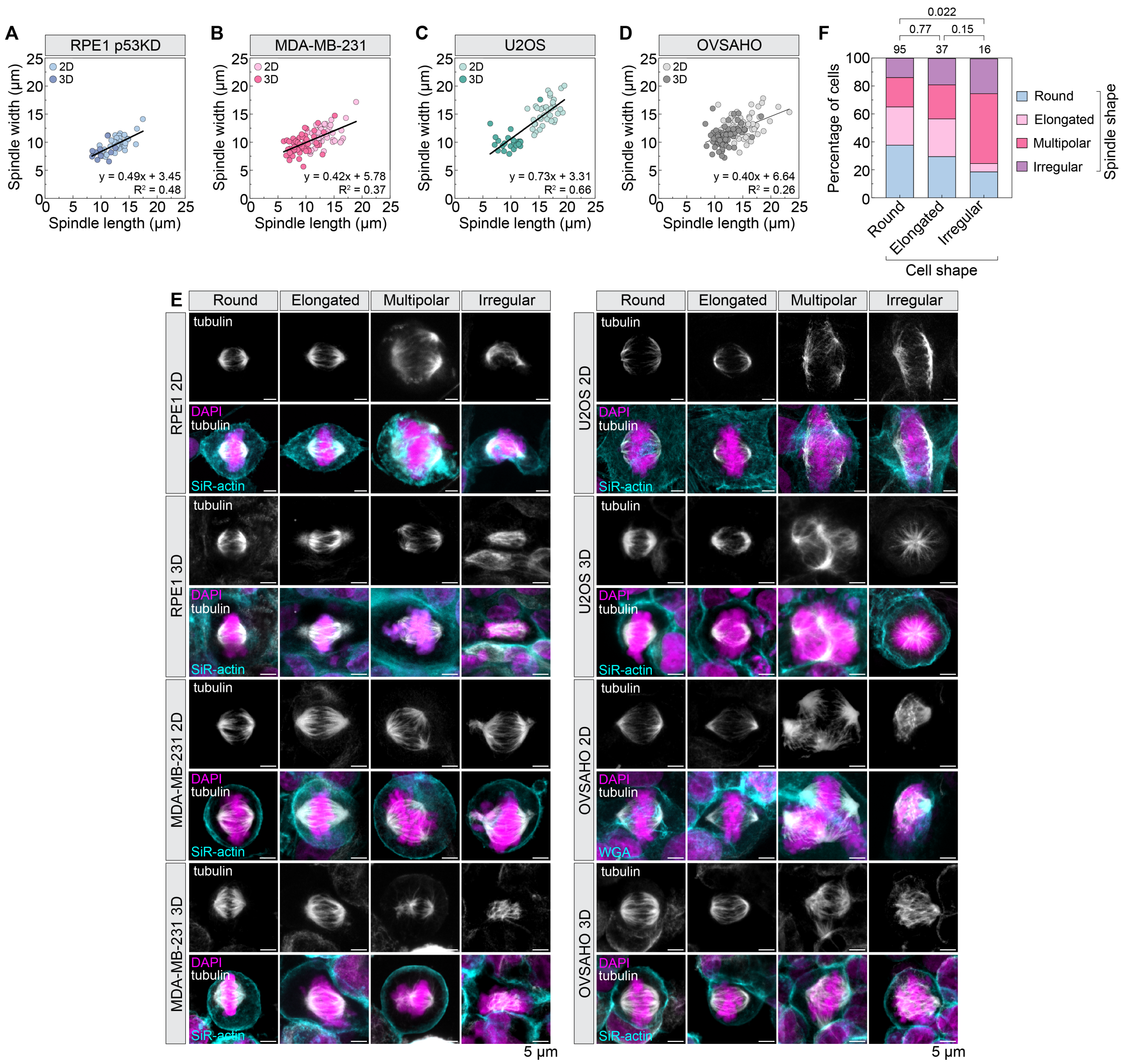


**Supplementary figure 3. Cells that compose spheroids possess smaller spindles whose irregular shape can be predetermined by the irregular cell morphology**

(A)-(D) Association between spindle length and spindle width in monolayers and spheroids stated for RPE1 p53KD (A), MDA-MB-231 (B), U2OS (C), and OVSAHO (D) cell lines. Each dot represents one cell.

(E) Examples of different spindle shapes and polarities for 4 different cell lines in 2D and 3D culture, as indicated. DAPI is used to label DNA (magenta), with immunostained tubulin (white). SiR-actin is used to stain the cell membrane (cyan) of all cell lines in both monolayers and spheroids, except for OVSAHO monolayers, where WGA is used to stain the actin cytoskeleton. The length of the scale bars is given in the images.

(F) The percentage of MDA-MB-231 cells with a particular shape that display a spindle phenotype as indicated in the legend.

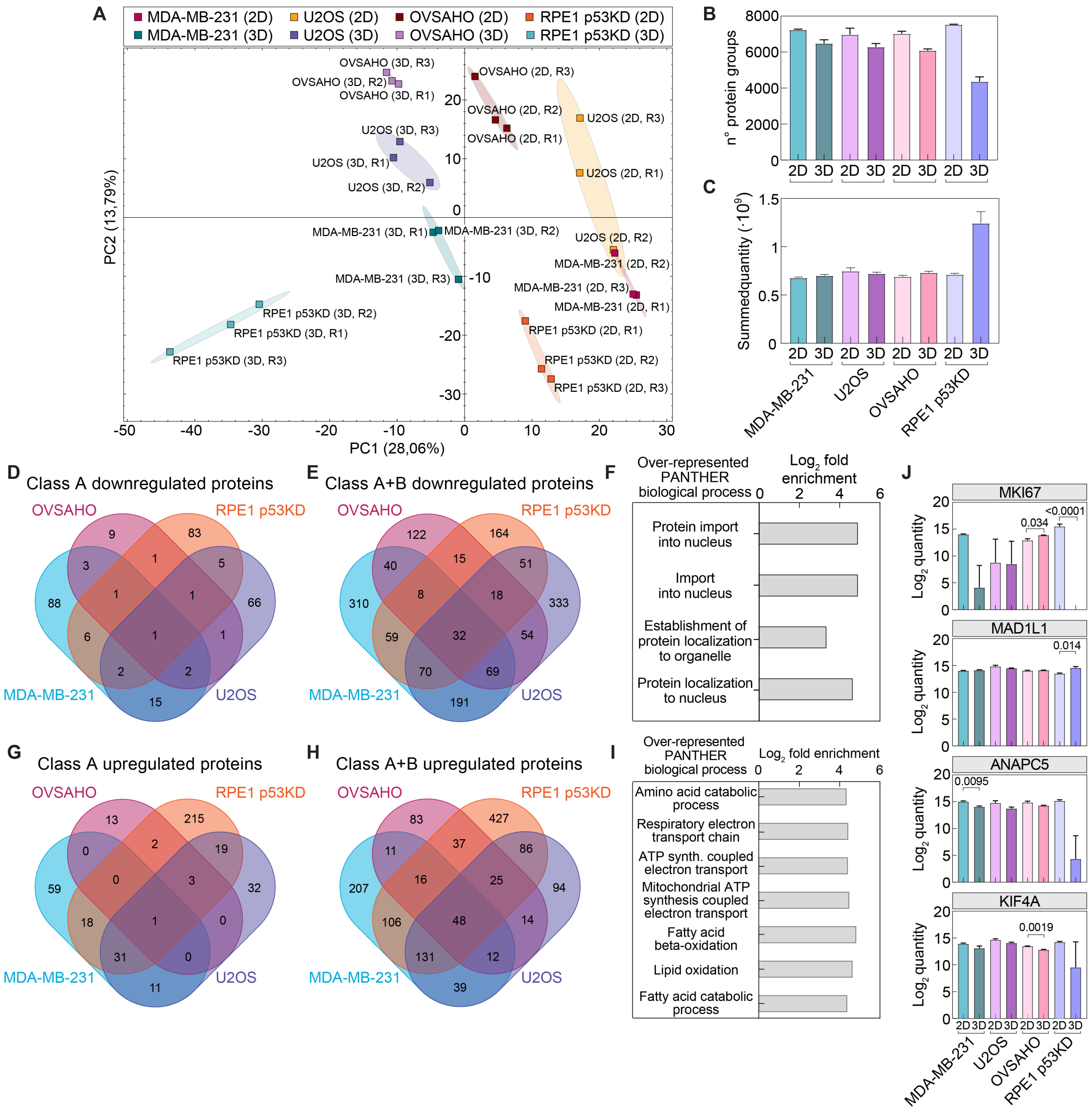


**Supplementary figure 4. Quality controls and additional data from proteome analysis**

(A) Principal component analysis shows clear separation of 2D and 3D cultures and cell lines and indicates the highest similarity between replicates.

(B) Total number of identified protein groups in 2D and 3D cultures of all cell lines. The mean of three technical replicates (n = 3) ± SEM is depicted.

(C) Summed quantity. The mean of three technical replicates (n = 3) ± SEM is depicted. Identified proteins plus the Venn diagram of all quantified proteins.

(D) Venn diagram of class A significantly downregulated proteins in 3D cultures.

(E) Venn diagram of class A and B significantly downregulated proteins in 3D cultures.

(F) Enrichment analysis of commonly significantly upregulated proteins in 2D cultures compared with 3D.

(G) Venn diagram of class A significantly upregulated proteins in 3D cultures.

(H) Venn diagram of class A and B significantly upregulated proteins in 3D cultures.

(I) Enrichment analysis of commonly significantly upregulated proteins in 3D cultures compared with 2D

(J) Log2 quantity of different proteins in 2D and 3D cultures across all cell lines. The mean of three technical replicates (n = 3) ± SEM is shown. A t-test was used to compare 2D versus 3D, with p-values stated above the bars.
